## Supplemental Figure 1 for "Flexible RSV prefusogenic fusion glycoprotein exposes multiple neutralizing epitopes that may collectively contribute to protective immunity"

**Appendix A**

***Method:*** *Competition of RSV F Site-specific mAb with Polyclonal Antibodies to RSV F Proteins by Bio-layer Interferometry (BLI)*

Competitive binding of antigenic site-specific mAb with serum antibodies from immunized cotton rats was determined by BLI using an Octet OK384 instrument (Pall FortéBio, Menlo Park, CA, USA). Amine-reactive biosensor tips (Ar2G) were activated with 20 nM N-(3-dimethylaminopropyl)-N’-ethylcarbodiimide (EDC) and 10 mM N-hydroxy succinimide ester (NHS) buffer for 300 s. 100 nM prefusion F(BV2129) was coupled to the amine-reactive tips in 10 mM acetate buffer (pH 6.0) for 600 s using the manufacturer’s protocol. Unreacted amine was quenched with 1M ethanolamine followed by washing to baseline for 300 s. Prefusion F (BV2129) coupled biosensor tips were allowed to associate with serially diluted serum antibodies for 600 s. Washed biosensor tips were then exposed for 400 s with competing mAbs D25 (site Ø), hRSV90 (site V), palivizumab (site II), R1.42 (site IV), or R7.10 (p27). Competition between serum antibodies and the RSV F mAbs was analyzed with OctetHT10.0 software. Competing antibody equivalent (CAE) levels (µg mL^-1^) were calculated based up on the percent competition, the concentration of competing antibody, and the serum dilution.

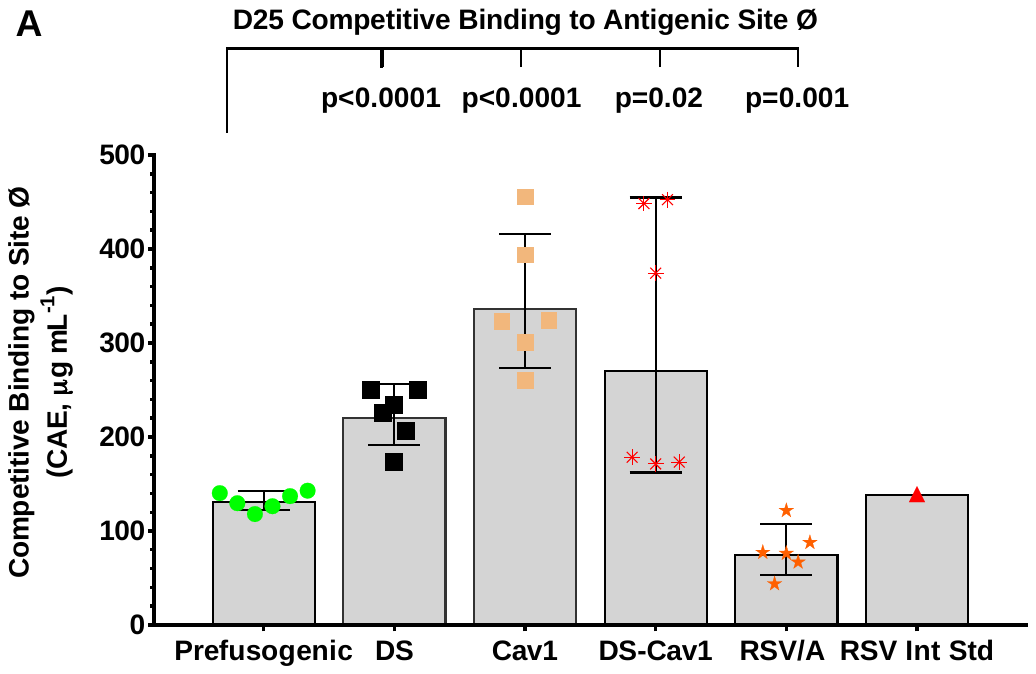

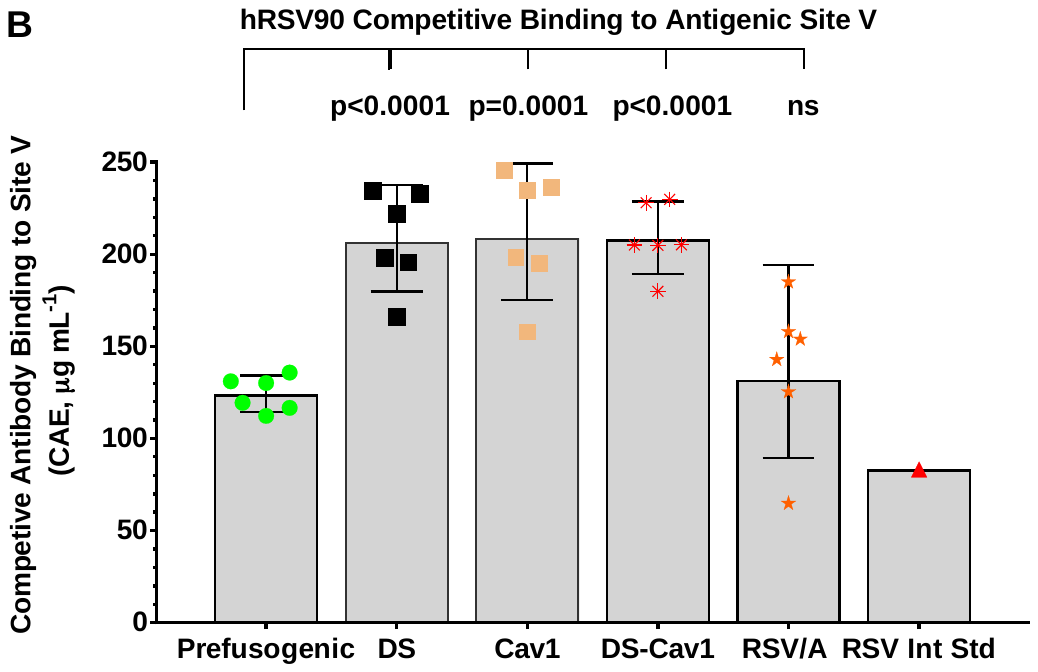

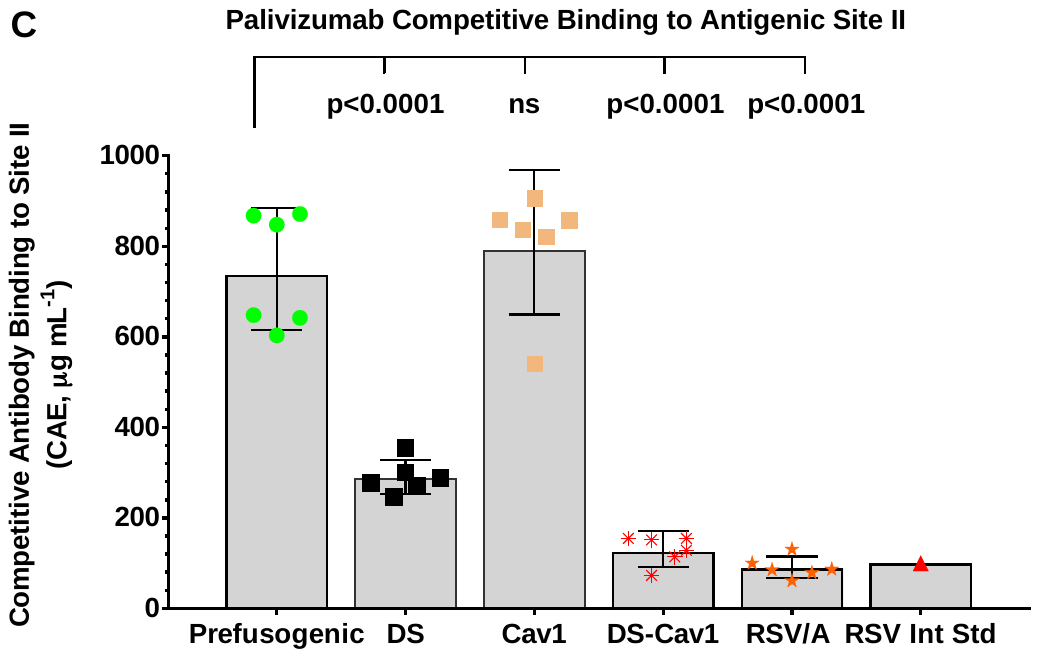

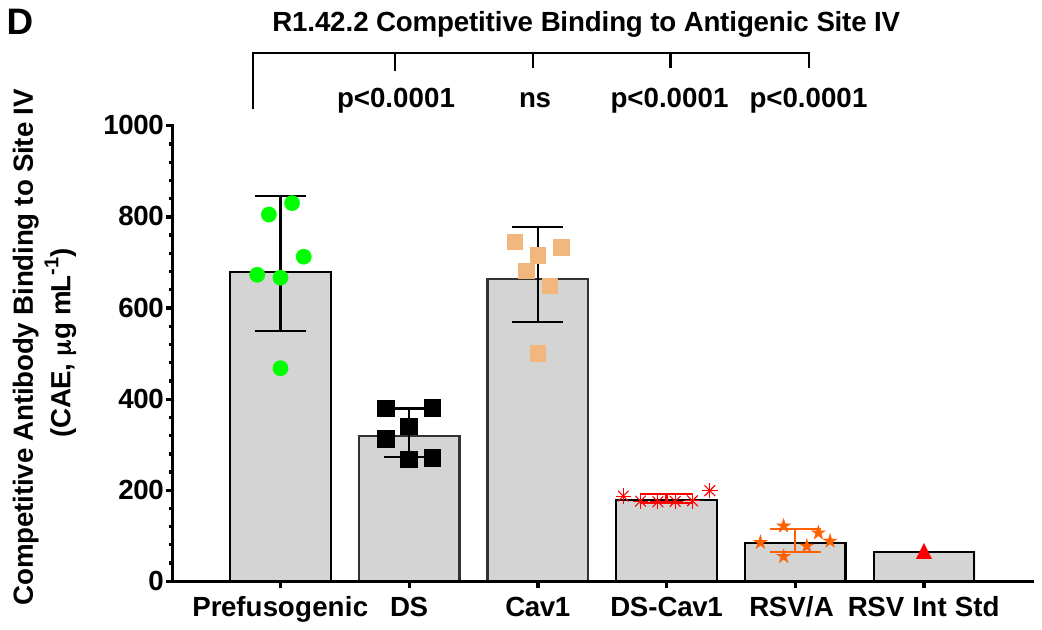

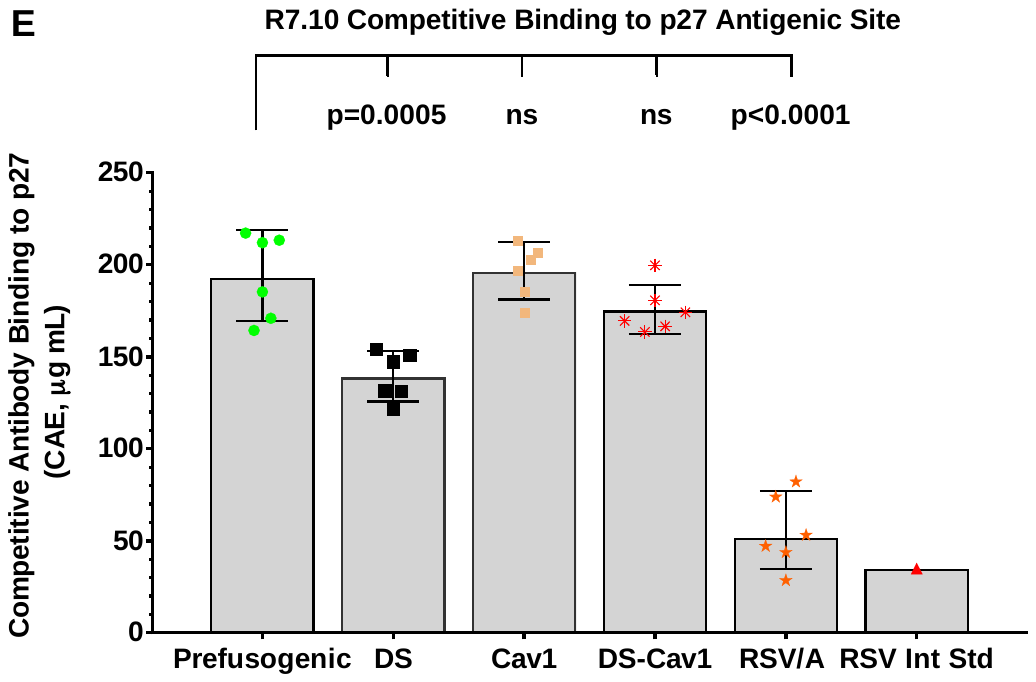

**Figure S1.** Competitive binding of RSV F site-specific monoclonal antibodies with immune serum determined by bio-layer interferometry (BLI) in cotton rats. Amine-reactive biosensor tips (ArG) were coupled to prefusion F (BV2129). Serum from cotton rats immunized with RSV F proteins was allowed to associate with biosensor tips for 600 s. Tips were washed and exposed for 400 s with site-specific competing mAbs. (**A**) D25 (site Ø). (**B**) hRSV90 (site V). (**C**) Palivizumab (site II). (**D**) R1.42 (site IV). (**E**) R7.10 (p27). Competition between serum antibodies and RSV F mAb was analyzed to OctetHT10.0 software. Horizontal bars indicate the group (N = 6 per group) GMT and the error bars indicate the 95% CI. Statistical significance between the RSV F prefusogenic vaccine group (BV1184) and paired immunized groups is indicated. Not significant (ns).
